# CAT Bridge: An Efficient Toolkit for Gene-Metabolite Association Mining from Multi-Omics Data

**DOI:** 10.1101/2024.01.21.576587

**Authors:** Bowen Yang, Tan Meng, Xinrui Wang, Jun Li, Shuang Zhao, Yingheng Wang, Shu Yi, Yi Zhou, Yi Zhang, Liang Li, Li Guo

## Abstract

**Background:** With advancements in sequencing and mass spectrometry technologies, multi-omics data can now be easily acquired for understanding complex biological systems. Nevertheless, substantial challenges remain in determining the association between gene-metabolite pairs due to the non-linear and multifactorial interactions within cellular networks. The complexity arises from the interplay of multiple genes and metabolites, often involving feedback loops and time-dependent regulatory mechanisms that are not easily captured by traditional analysis methods.

**Findings:** Here, we introduce Compounds And Transcripts Bridge (abbreviated as CAT Bridge, available at https://catbridge.work), a free user-friendly platform for longitudinal multi-omics analysis to efficiently identify transcripts associated with metabolites using time-series omics data. To evaluate the association of gene-metabolite pairs, CAT Bridge is a pioneering work benchmarking a set of statistical methods spanning causality estimation and correlation coefficient calculation for multi-omics analysis. Additionally, CAT Bridge features an artificial intelligence (AI) agent to assist users interpreting the association results.

**Conclusions:** We applied CAT Bridge to experimentally obtained *Capsicum chinense* (chili pepper) and public human and *Escherichia coli* (*E. coli*) time-series transcriptome and metabolome datasets. CAT Bridge successfully identified genes involved in the biosynthesis of capsaicin in *C. chinense*. Furthermore, case study results showed that the convergent cross mapping (CCM) method outperforms traditional approaches in longitudinal multi-omics analyses. CAT Bridge simplifies access to various established methods for longitudinal multi-omics analysis, and enables researchers to swiftly identify associated gene-metabolite pairs for further validation.

## Background

Recent advancements in sequencing and mass spectrometry (MS) technologies have made the acquisition of multi-omics data more cost-efficient and feasible, multi-omics data analysis is crucial for understanding intricate biological mechanisms from a more comprehensive perspective than single omics, this holistic analysis allows us to explore the interplay between different molecular levels [1–4]. For integrated data analysis of transcriptomics and metabolomics, a crucial task is to examine the associated gene-metabolite pairs. Existing strategies bifurcate primarily into two classes: knowledge-driven approaches and data-driven approaches [5]. Knowledge-driven approaches have shown their inadequacies for non-model organisms due to the lack of knowledge, restrictions in revealing *de novo* mechanisms, and difficulties in quantifying and ranking their outcomes [5]. Data-driven strategies mainly depend on statistical methods that model the correlation of gene-metabolite pairs or sophisticated machine learning methods [6, 7]. However, due to the severe batch effects in omics data, machine learning approaches usually lack generalizability [8]. Meanwhile, they are also prone to overfit when applied to relatively small omics datasets, making them harder to transfer to different scenarios and less interpretable [5]. In terms of statistical methods, people usually calculate the correlation coefficient to match gene-compound pairs [9, 10] such as in the studies of the growth cycles of *Solanum lycopersicum* (tomato) [11] and *Oryza sativa* (rice) [12], where Pearson correlations were utilized to study metabolic regulatory networks by integrating transcriptomics and metabolomics data. However, these methods face reliability issues, especially when dealing with longitudinal omics data. This is because of the time lag in the expression of genes and metabolites, and the inherent complexity of biological systems, which is a dynamical system with non-linear interactions between different molecules [4, 13]. A more reliable solution is to use causality, which is inferred based on the ability of one time series (e.g., gene expression) to predict another (e.g., metabolite concentration), to replace the correlation coefficient [14, 15]. Furthermore, purely data-driven strategies can occasionally lead to biologically naive conclusions [9]. For example, a purely data-driven approach might incorrectly link low cholesterol levels with higher mortality rates, suggesting that lower cholesterol is detrimental. However, the actual cause is underlying diseases like cancer, which cause both low cholesterol and increased mortality [16]. Therefore, integrating data-driven and knowledge-driven methodologies may offer a more comprehensive and accurate interpretation of multi-omics data.

To address the existing limitations, we have introduced Compounds And Transcripts Bridge (CAT Bridge), a comprehensive cross-platform toolkit that provides a novel analysis pipeline for integrative analysis linking upstream and downstream omics (typically transcriptomics and metabolomics). The novel pipeline encompasses three essential steps, data preprocessing, computing association between gene-metabolite pairs, and result presentation. For measuring the association, we integrated seven different statistical algorithms on causality estimation and correlation coefficient calculation and benchmarked them on human, plant, and microorganism datasets. It also offers three ways to display results that are generated from both data-driven approach and knowledge-driven approach, including common omics statistical analysis and visualization, heuristic ranking of candidate genes based on causality or correlation, and an AI agent driven by large language models (LLMs) to identify associated gene-metabolite pairs through prior knowledge **(Figure 1)**.

**Figure 1.**
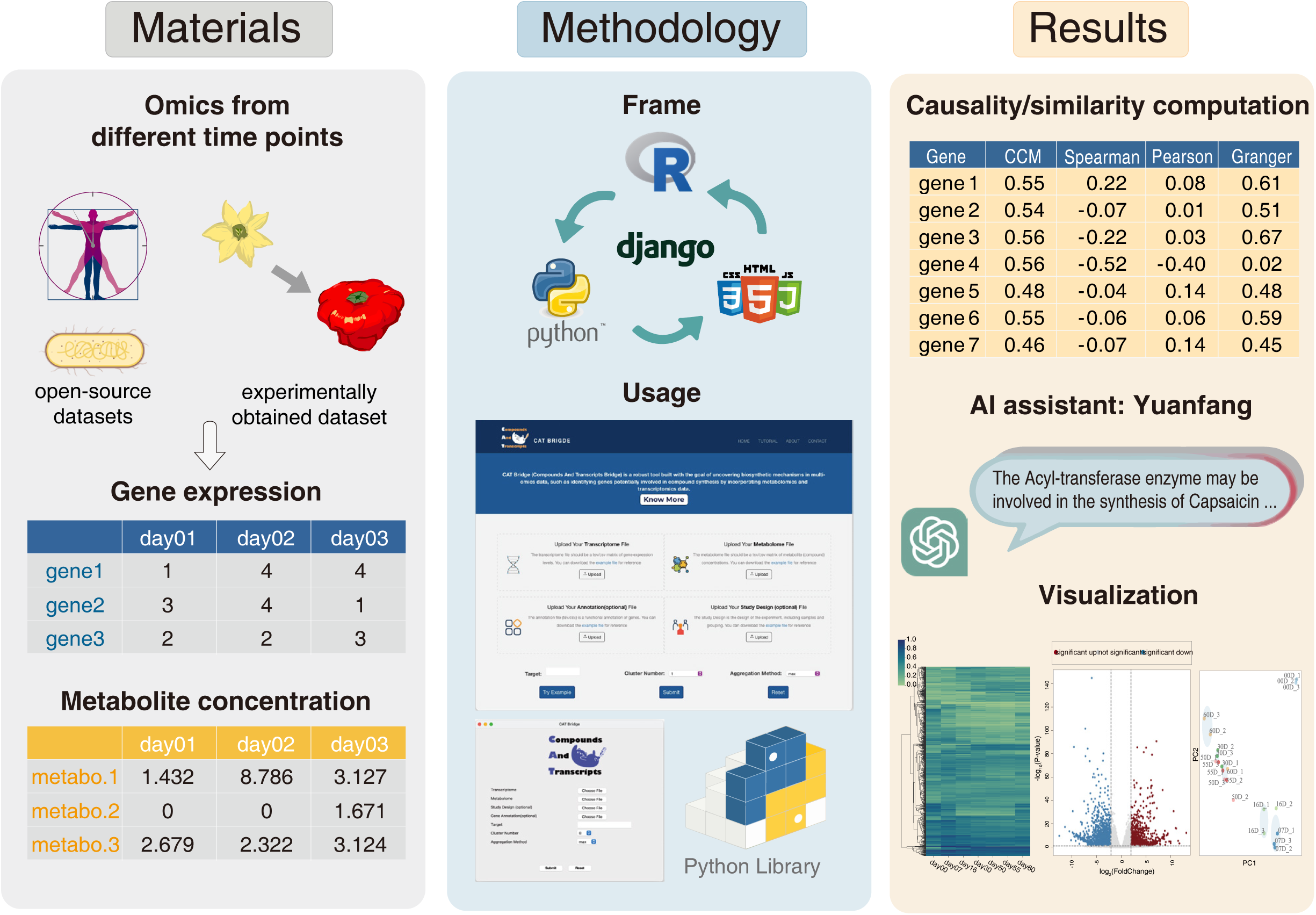
The architecture of CAT Bridge project. Benchmark data come from publicly available datasets and our experimentally obtained *C. chinense* dataset, including gene expression and metabolite concentration matrices. The construction of CAT Bridge relies on Python, R, HTML/CSS, and JavaScript, and provides three modes of usage: web server, standalone program, and Python library. To assist in the discovery of associated gene-metabolite pairs, it provides results from three perspectives: statistical analysis, AI agent-generated responses, and data visualization.

Besides, we offer three different access options for CAT Bridge, including a web server, a standalone application, and a Python library. We also provide a detailed tutorial and a sample dataset to help the users get started easily.

## Materials and Methods

### Overview of CAT Bridge

The workflow of CAT Bridge consists of three primary steps, (1) data processing; (2) statistical modeling (causality estimation and correlation coefficient calculation); (3) visualization and interpretation (**Figure 2A**). Users are required to upload two processed files, gene expression and metabolite concentration matrices, and specify a metabolite of interest as the target. After data processing, seven different causality and correlation algorithms are applied to measure the relationship between each gene and the target metabolite. The algorithm chosen by the user will then generate a vector for each gene representing one-to-one associations between the gene and the target metabolite (**Figure 2B**). Subsequently, a vector magnitude is employed for heuristic ranking, with the top 100 genes being reviewed by the AI agent, to utilize prior knowledge to offer inspiration to users. Finally, commonly used omics visualization will be employed to assist users in interpreting the overall expression pattern and facilitating the selection of potential candidate genes.

**Figure 2.**
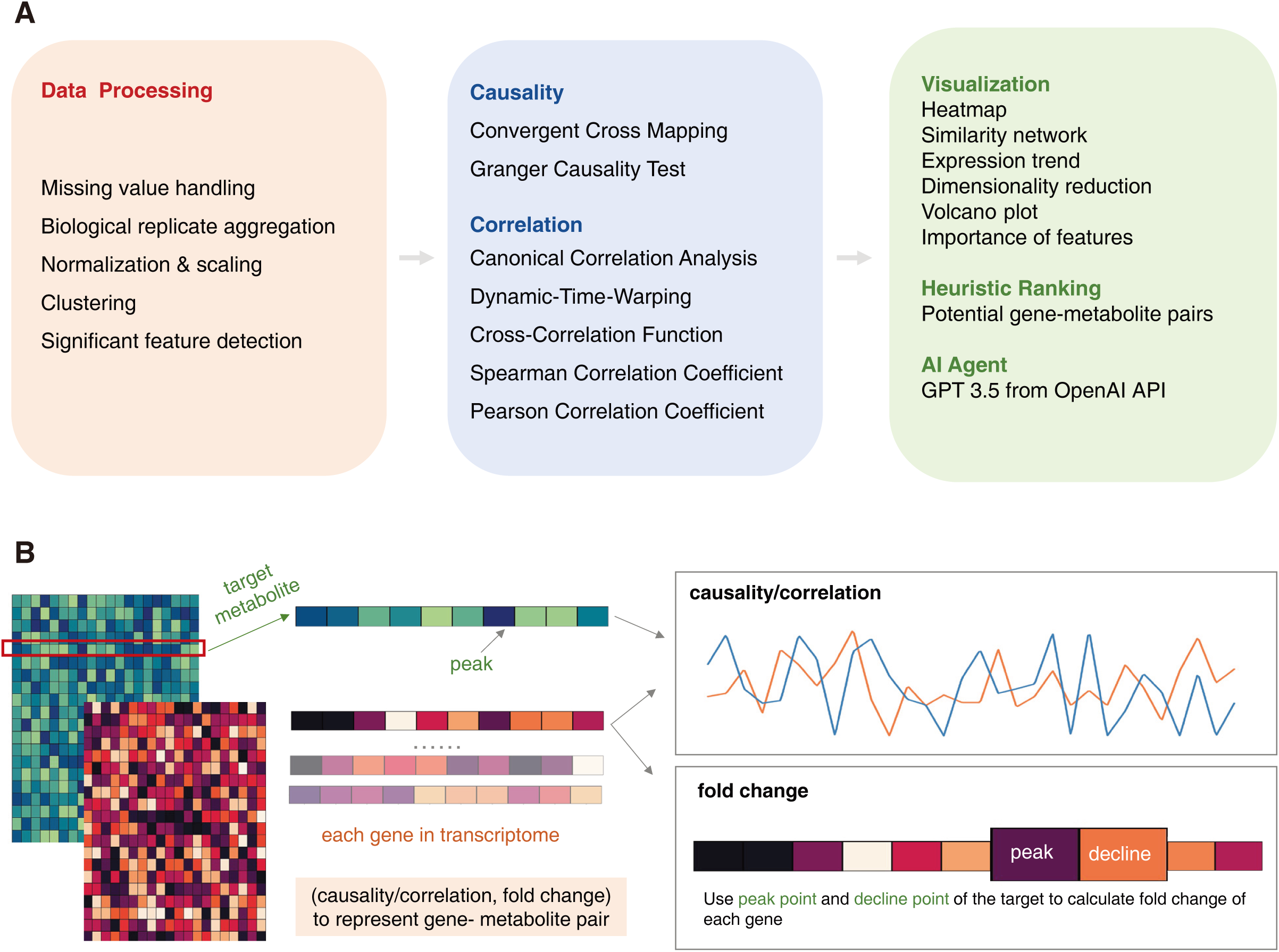
The features and overall workflow of CAT Bridge. (A) The workflow of CAT Bridge consists of three primary steps: data preprocessing, estimation of cause-effect relationships or computation of correlation coefficient, and the presentation of results, which includes visualization, heuristic ranking, and responses from an AI agent. (B) The computation of CAT Bridge involves: extracting the target metabolite from the metabolite concentration matrix, and pinpointing the time point of its maximum concentration as the peak time. Next, the causality or correlation between each gene in the gene expression matrix and the target metabolite is obtained. Then along with the fold change between the peak and decline time points to compose a vector to represent the association between gene-metabolite pairs.

### Gene-metabolite association computing

For gene-metabolite pair identification such as inferring the biosynthetic genes for particular metabolites, correlation is often used to imply association, such as Spearman Correlation Coefficient (Spearman) and Pearson Correlation Coefficient (Pearson). However, such correlation-based methods have substantial limitations [9, 10] because they overlook the non-linearity and lag issues of gene expression leading to metabolite changes. Therefore, besides considering Spearman and Pearson, we have integrated into CAT Bridge various distinct statistical methods, including Granger Causality (Granger) and Convergent Cross Mapping (CCM) by evaluating the predictability of metabolite concentration from gene expression to represent causality, as well as Canonical Correlation Analysis (CCA), Dynamic-Time-Warping (DTW), Cross-Correlation Function (CCF) for calculating correlation coefficient (The implementation methods are provided in the Supplementary Text). These algorithms were based on different assumptions so that some of them allow compatibility with time series data and complex systems (**Table 1**). Among them, correlation-based strategies have been widely applied in genomics and multi-omics analysis [17–20]. The CCM and Granger, which estimate causality from time series data, are already used in some areas of biology such as ecology and neurobiology, but are overlooked in the omics analysis [14, 21–23]. Furthermore, our benchmarking results (as detailed in the Results section) suggest that in longitudinal multi-omics studies, causal relationships may provide a more accurate depiction of the association between genes and metabolites.

**Table 1.** Comparison of statistical methods by linearity and time lag consideration.

| Statistical Method | Linearity | Time Lag Consideration |
| --- | --- | --- |
| Pearson Correlation Coefficient (Pearson) | Non-linear | No |
| Spearman Correlation Coefficient (Spearman) | Non-linear | No |
| Convergent Cross Mapping (CCM) | Non-linear | Yes |
| Granger Causality Test (Granger) | Linear | Yes |
| Canonical Correlation Analysis (CCA) | Linear | No |
| Dynamic-Time-Warping (DTW) | Non-linear | Yes |
| Cross-Correlation Function (CCF) | Linear | Yes |

### Heuristic ranking of candidate genes

Fold change (FC) is another measurement frequently used in omics analyses to identify differentially expressed genes [24]. CAT Bridge pinpoints the peak time of the target metabolite and calculates each gene’s log2 normalized FC of this peak time point and the subsequent decline time point (that is, the next sampling time point after the peak time point) using DESeq2 [25]. Then, causality or correlation and FC are combined into a vector to represent the gene-metabolite pair. After the min-max normalization of values (The details are provided in the Supplementary Text), the magnitude of this vector is calculated as the CAT score (Figure 2B). This score heuristically ranks the strength of association between each gene and the metabolite. Users can filter putative genes based on thresholds (e.g., 0.5 for causality and correlation, 1 for normalized FC) or manually review them in descending order.

### Knowledge-driven approaches and visualization

Optionally, if users provide a gene function annotations file, typically derived from homology annotations using tools like InterProScan [26] or eggNOG-mapper [27] for non-model organisms), the CAT score will be adjusted with an additional value. This value is determined by a scoring rule based on the gene’s description. By default, genes annotated as enzymes receive a score of 0.2, while those with unknown functions get a score of 0.1. Users can customize this scoring rule based on their specific requirements, depending on the presence of target annotations and their importance. Finally, the top 100 genes in heuristically ranking will be evaluated by the GPT-3.5 Turbo based AI agent, to identify putative genes on the gene’s functional annotation and prior knowledge (The implementation methods are provided in the Supplementary Text). To enhance data interpretation, the CAT Bridge workflow offers a visual ranking of genes based on computation results, and also incorporates a spectrum of widely utilized graphical outputs, such as heatmap, and principal component analysis (PCA) plot, details can be found in the Supplementary Text.

### Plant Material Cultivation and Sampling

To test the effectiveness of CAT Bridge across different species, especially its applicability to non-model organisms, we collected transcriptome sequencing and metabolic profiling data from *C. chinense*, focusing on one of its trademark natural products, capsaicin. *C. chinense* seedlings were divided into three groups, each containing 15 seedlings, grown in a greenhouse of Peking University Institute of Advanced Agricultural Sciences with a controlled environment of 25°C temperature, a light-dark cycle of 16 hours light and 8 hours dark, and 70% relative humidity. The fruits of *C. chinense* were sampled at seven distinct time points, starting from the day of flowering, i.e. 0 day post-anthesis (DPA) during which flowers were collected. Followed by fruit harvest on days 7, 16, 30, 50, 55, and 60 DPA. For each time point, we sampled 15 fruits from each group of seedlings, yielding a total of 45 sampling per time point. For each group, we utilized 1.0 mL of 70% aqueous methanol per sample, with a sample weight of approximately 50 mg (aside from the 7 DPA samples which averaged 27.3mg). The samples were ground and freeze-dried in liquid nitrogen. Each sample was then extracted using 1.0 mL of 70% aqueous methanol for every 50 mg of sample. Following extraction, the samples underwent ultrasonic treatment using an Ultrasonic Cell Disruptor SCIENTZ-IID (Ningbo Scientz Biotechnology Co., LTD., China) at a frequency of 40 kHz for 10 minutes at room temperature. Standards were prepared as follows: a mixed standard solution, ranging from 20-50 μg/mL, was prepared using MS-grade methanol. For the amino acid standard solution, a 1 mg/mL stock solution was initially prepared in water, and then diluted with 50% methanol to achieve a final concentration of 50 μg/mL. Three biological sample replicates were utilized in the subsequent transcriptome and metabolome analyses.

### Metabolome Profiling using HPLC-MS and Data Pre-processing

The metabolome profiling was carried out using untargeted metabolomics based on liquid chromatography coupled with mass spectrometry (LC-MS). The samples were filtered through a 0.22 µm membrane and transferred into the lining tube of a sampling vial. Subsequent centrifugation was carried out at 12000 rcf and 4℃ for 10 minutes. The processed samples were then analyzed using Thermo Scientific Orbitrap Exploris™ 240 (Thermo Fisher Scientific, USA). Chromatographic separation was achieved on a T3 C18 (1.7 µm, 2.1 mm × 150 mm column, USA) maintained at 40℃. The mobile phase consisted of A: 1% formic acid in water and B: 1% formic acid in acetonitrile, with a flow rate of 300 µL/min. A 3 µL sample was injected at an autosampler temperature of 10℃. The elution gradient was set as follows: 0-2.5 min, 3-10% B; 2.5-6 min, 10-44% B; 6-14 min, 44-80% B; 14-20 min, 80-95% B; 20-23 min, 95% B; 23-23.1 min, 95-3% B; 23.1-28 min, 3% B. MS was performed using both positive and negative ion scans, with a precursor ion scan mode. The auxiliary gas heater temperature was set at 350℃, and the ion transfer tube temperature was also maintained at 350℃. The sheath gas flow rate and auxiliary gas flow rate were set to 35 arb and 15 arb, respectively. The voltages were set to 3.5 KV for the positive spectrum and 3.2 KV for the negative spectrum. For MS1, the scan resolution was 60000, with a scan range of 80-1200. For MS2, the scan resolution was 15000, with a stepped collision energy of 20, 40, and 60 eV. Metabolite identification and quantification were performed using the Compound Discoverer software 3.3 (Thermo Fisher Scientific, USA).

### RNA extraction and transcriptome sequencing

Total RNA was isolated from the above collected plant materials using Trizol Reagent (Thermo Fisher, USA) following manufacturer recommended protocol. The quality of RNA extracts was evaluated using RNA Nano 6000 Assay Kit of the Bioanalyzer 2100 system (Agilent Technologies, USA) following manufacturer’s recommendation and samples with a RIN value >7 were used in downstream sequencing library construction and sequencing. The library construction was conducted using Illumina True-seq transcriptome kit (Illumina, USA) following standard protocols. Transcriptome sequencing was carried out by Novogene Co., Ltd. The sequencing reads were procured from the Illumina NovaSeq 6000 platform. For pre-processing, fastp [28] was employed to conduct quality control and clean the data. Subsequently, these reads were mapped to the *C. chinense* cultivar PI159236 genome [29] using STAR [30]. StringTie [31] was utilized to quantify and assess the expression levels of the genes that were successfully mapped. The biological function annotation of genes is obtained through the eggNOG-mapper.

### Acquisition and processing of public datasets

We also collected human and *Escherichia coli* (*E. coli*) multi-omics data from public datasets to further examine the performance of CAT Bridge.

The human data was sourced from a published aging study [32] that sampled transcriptome and metabolome data from 28 younger (20 to 25 years) and 54 older (55 to 66 years) female human donors. 13 time points are included in them where both the transcriptome and the metabolome data were detected. We obtained the transcripts expression levels and the concentrations of glucose and fructose 6-phosphate for 13 donors, if multiple donor data were available at a single time point, the average value was used.

The dataset for *E. coli* originated from a study on the interactions between metabolites and genes within the bacterium [33]. Researchers manipulated the culture conditions of *E. coli*, alternating between growth and starvation phases. They then collected the transcriptome and metabolome under these varying conditions to systematically explore how metabolites influence transcription factors. The dataset includes 29 time points where both transcriptomic and metabolomic information were concurrently obtained.

## Results

CAT Bridge provides a platform with a novel pipeline that allows for the rapid identification of putative genes for further investigation and validation. And three distinct usage modes are offered to cater to a wide range of user requirements. Firstly, it features a web server, designed for user-friendliness and accessibility, and is open to all users without any login requirements. This is particularly beneficial to those who are less familiar with programming languages. Secondly, a standalone application is available for users handling large data files, as it lifts the constraints on file sizes. Finally, a Python library is available for bioinformaticians with complete features and customizable workflows (The implementation of the software is provided in the Supplementary Text).

To showcase the utility and features of CAT Bridge, we applied it to both an experimentally obtained dataset collected from *C. chinense* and a publicly available human and *E. coli* dataset in two case studies. Our analysis revealed that in the context of longitudinal multi-omics, causality-based strategies tend to outperform those solely based on similarity. As such, we advocate for the adoption of causality rather than similarity in longitudinal multi-omics analysis such as co-mining the transcriptomics and metabolomics data. The capsaicin dataset generated in this study has been made open source for user exploration.

### Case Study 1: Identifying genes associated with capsaicin biosynthesis in *C. chinense*

In our inaugural case study, we leveraged experimentally obtained non-model organism data to examine the performance of CAT Bridge. This data comprised the transcriptome and metabolome of *C. chinense* at seven different developmental stages after bloom (**Figure 3A**). Capsaicin, an important natural product produced by *C. chinense* that gives fruit pungency and has potential anti-cancer and analgesic activity [34], was selected as the target metabolite for this study. Time-series transcriptome and metabolic profiling of developing *C. chinense* fruits were used as input data to test CAT Bridge.

**Figure 3.**
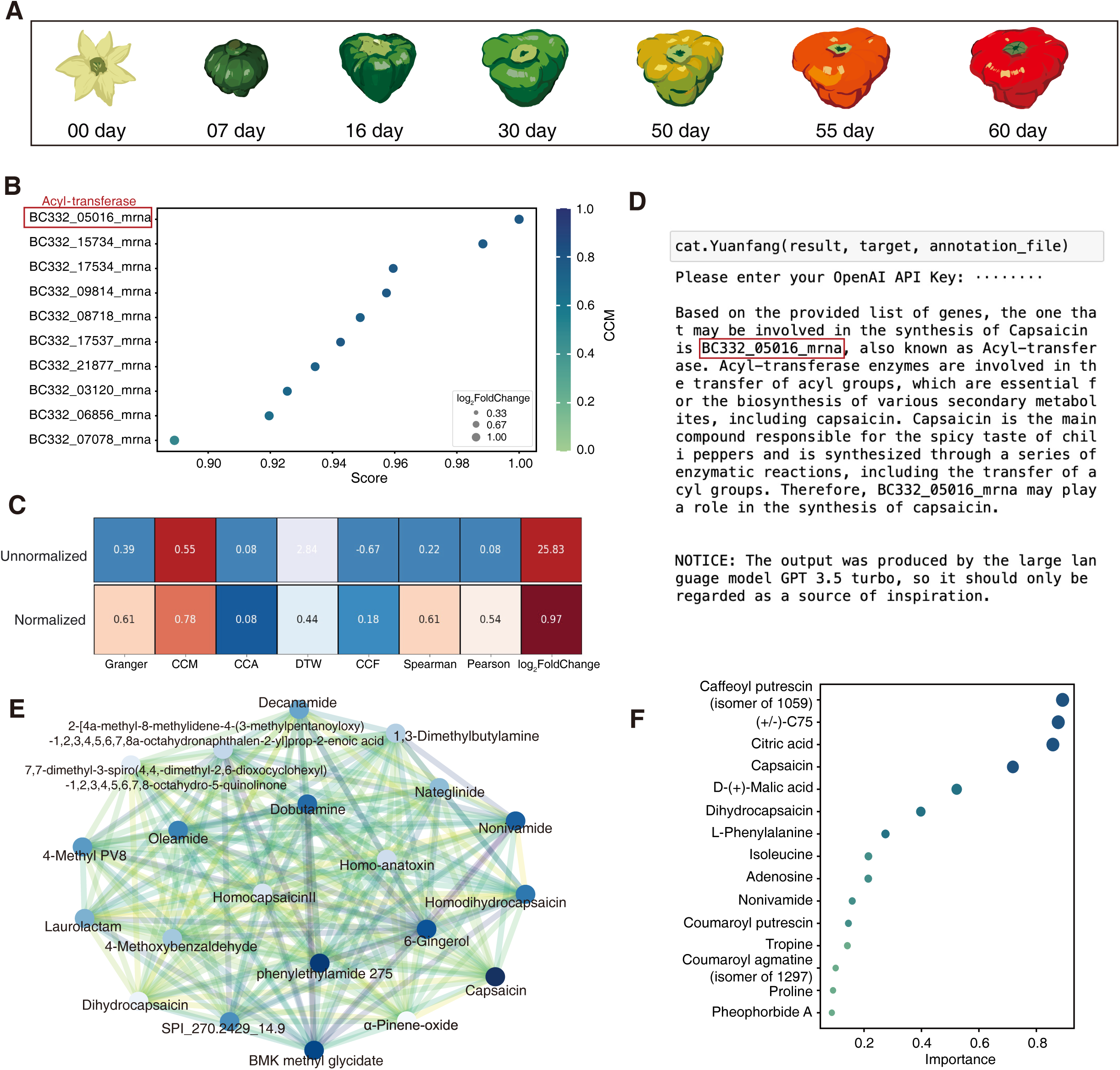
Application of CAT Bridge to mine transcript-capsaicin association. (A) Diagram showing the time points sampled for transcriptome and metabolic profiling during fruit development of *C. chinense* in case study 1. (B) Heuristic ranking produced using the CCM-based method. (C) Comparative values across different methods. For unnormalized values: red indicates a strong association; original denotes medium association; blue suggests values that are below the commonly used threshold, show no association, or are negatively associated (depending on the method); light blue means this method does not adhere to a common threshold. For normalized values: red signifies values that are high after min-max normalization; blue represents low normalized values. (D) Interpretation results derived from the AI agent (E) The correlation network of capsaicin. (F) The significance of metabolites.

Through examination using the CCM method for hypothetical ranking, BC332_05016 encoding an Acyl-transferase was ranked first, regardless of whether an annotation file was provided. The Result suggests that this gene was more likely to be the synthetic gene associated with capsaicin in *C. chinense* (**Figure 3B**). The complete heuristic ranking results are provided in Supplementary Table S1. BLAST search revealed that BC332_05016 was homologous of PUN1 (sequence identity: 100%), a.k.a AT3 (Acyl-transferase 3) or CS (capsaicin synthesis) gene [35]. Moreover, when common thresholds were applied for screening, only CCM passed the criteria. The causality modeled based on CCM was 0.55, implying a strong association between BC332_05016 and capsaicin. By contrast, the conventional Pearson correlation method produced a result of 0.08, which would fall below the commonly used threshold and potentially lead to an overlook of this gene-metabolite pair (**Figure 3C**). The AI agent also accurately found BC332_05016 among the top 100 genes based on functional annotation (**Figure 3D**). Furthermore, CAT Bridge visualization result showed that capsaicinoids such as nonivamide, dihydrocapsaicin, and homocapsaicin have high similarity to capsaicin (**Figure 3E**) and may play a significant role in response to the variable (**Figure 3F**). The rest of the visualization results are provided in the Supplementary Figures (**Figure S1-S4**). These results show that the CAT Bridge is a valuable tool in multi-omics analysis to reliably identify associated gene-metabolite pairs.

### Case Study 2: Identifying association of gene-metabolite pairs in glycolysis

For case study 2, we utilized an open-source multi-omics dataset generated by previous aging research (Kuehne et al. 2017) [32]. After processing, 13 time points from the skin of younger and older people used for testing, we particularly compared different association modeling results from various statistical methods in CAT Bridge.

Using this dataset, Kuehne et al. found glycolysis altered activity in the upper body when aging, and hexokinase 2 (HK2) and phosphofructokinase (PFKP) two enzyme genes are reduced, whereas fructose bisphosphatase 1 (FBP1) and aldolase A (ALDOA) are increased in the older skin [32]. Because glycolysis is well-researched, and the study’s focus was not identifying gene-metabolite pairs, they only compared the fold changes in genes and metabolites, without evaluating the strength of associations between gene-metabolite pairs.

To explore whether different statistical methods can identify known gene-metabolite pairs in glycolysis, we used this dataset to evaluate the relationship between key genes and glucose, fructose 6-phosphate, for fold change, we used the same comparison methods as Kuehne et al., that is older group divided by the younger group. The results show that CCM still performed the best, accurately identifying a strong negative correlation between the concentration of glucose and the expression of HK2; a strong positive correlation between fructose 1,6-bisphosphate and PFKP, and a weak negative correlation with ALDOA. However, only Spearman yielded the expected relationship between fructose 1,6-bisphosphate and FBP1 (**Figure 4A, B**). Additionally, previous studies did not identify hexokinase 1 (HK1) as a significant gene in the glucose change because its levels in the elderly were slightly higher than in younger individuals. However, CCM correctly identified this relationship. Taking the expression pattern of PFKP and fructose 1,6-bisphosphate as an example, we observed that changes in gene expression were always reflected in the metabolite concentration at the next time point, indicating a delay between metabolite and transcript responses. This illustrates the advantage of causal relationship modeling methods over traditional methods.

**Figure 4.**
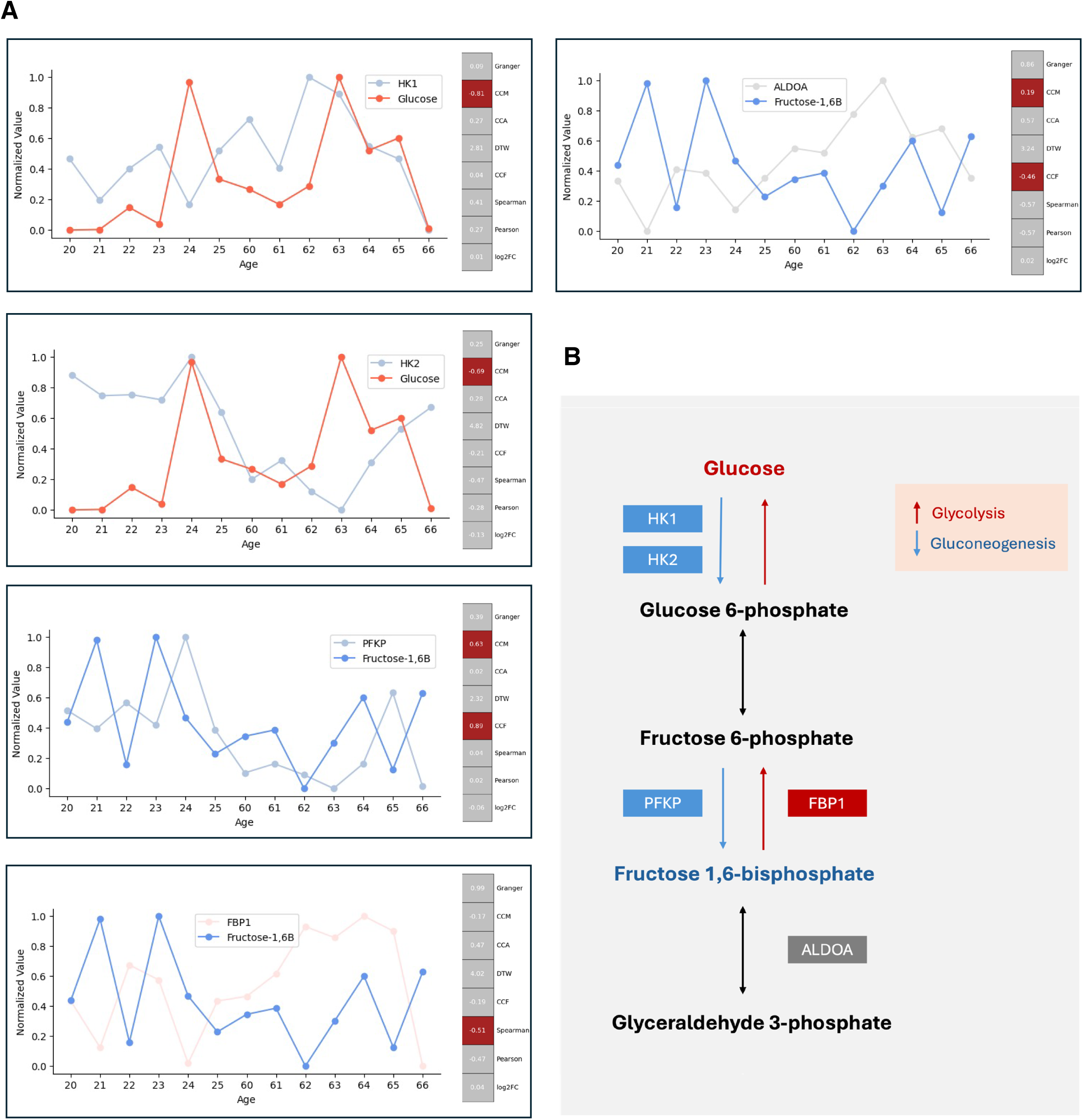
Application of CAT Bridge to mine transcript-metabolite associations in glycolysis pathway. (A) Left side: the expression patterns of functional genes and their corresponding target metabolites. Right side: the strength of associations was evaluated using different methods, red denotes evaluation outcomes that align with known pathway information under typical thresholds (e.g., a correlation coefficient < −0.5 indicating a strong negative association), whereas grey signifies inconsistent or unsuitable. (B) Part of the glycolysis pathway.

### Case Study 3: metabolite-transcription factor interactions and Acetyl-CoA regulation

In the third study, we utilized a transcriptomic and metabolomic dataset from *E. coli* to demonstrate the capabilities of CAT Bridge in analyzing simple organisms. The dataset contains gene expression and metabolite concentration data across 29 time points, and it was originally generated by Lempp et al. to investigate the allosteric regulation of transcription factors (TFs) by metabolites [33]. They constructed a “literature network” of known metabolite-TF interactions, including 16 interactions as activations of the TF by metabolites. Using Kinetic Correlation, which accommodates time lags, Lempp et al. recovered 10 out of the 16 activation interactions. We re-analyzed this dataset using CAT Bridge to test its performance with simple organisms. Utilizing the functions integrated within CAT Bridge and setting a time lag of 4, we calculated the association strengths between these molecules. The CCM method once again performed best, reproducing 14 out of 16 interactions.

Focusing on Acetyl-CoA as the target metabolite and setting a time lag of 3, we investigated the associated genes. The CCM-based methods fared the best, and the results showed strong causality (greater than 0.9) with the aceE, aceF, and lpd genes, which are involved in the conversion of pyruvate to Acetyl-CoA (**Figure 5A**) [36]. These genes were also ranked in the top 100 in heuristic sorting and clustered together. Additionally, the acs and pta genes, which are involved in the conversion of acetate to Acetyl-CoA, [36], also showed strong causality (0.840 and 0.721, respectively) but did not rank in the top 100 in heuristic sorting (**Figure 5B**). The complete heuristic ranking results are provided in Supplementary Table S2. This suggests that in this study, the precursors for Acetyl-CoA may predominantly come from pyruvate.

**Figure 5.**
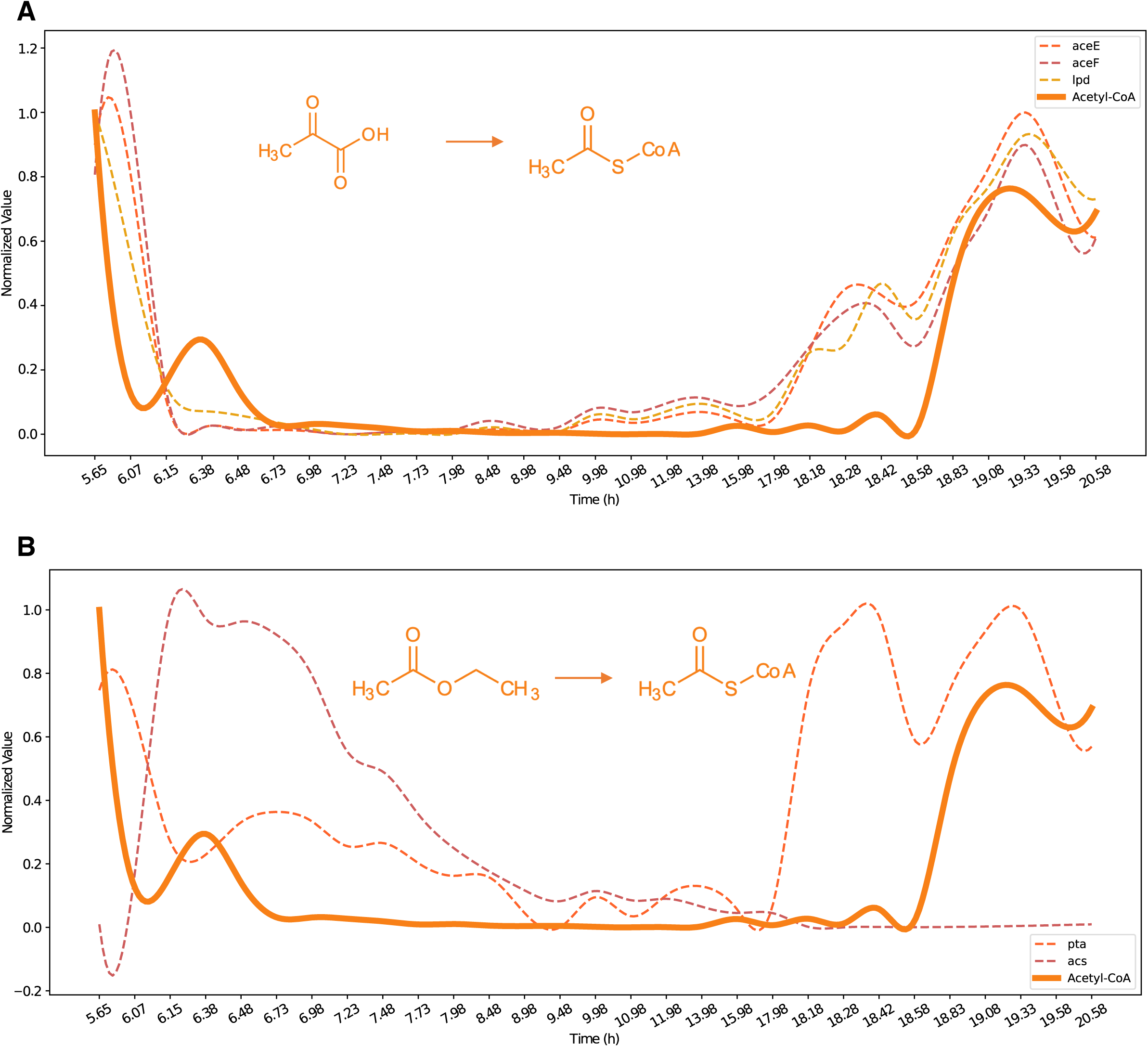
Expression Patterns of Acetyl-CoA Synthesis Genes and Acetyl-CoA Concentration. (A) Genes involved in the conversion of pyruvate to Acetyl-CoA. (B) Genes involved in the conversion of acetate to Acetyl-CoA.

These results demonstrate the CAT Bridge’s potential to extract meaningful insights from multi-omics data across diverse species. By integrating time-series analysis methods, particularly CCM, it offers superior performance in longitudinal omics compared to common methods.

### Comparison with other web-based tools

**Table 2** displays the function coverage comparisons between CAT Bridge and other data-driven multi-omics analysis web-based tools, including OmicsAnalyst [5], 3omics [37], IntLIM [38, 39], and CorDiffViz [40]. In association identification, IntLIM, 3omics, and CorDiffViz integrate either Pearson or Spearman correlations, or both, to aid in the discovery of feature relationships. What sets CAT Bridge apart is its assembly of various algorithms that handle time-series data and causality, and incorporate an AI agent to inspire users. Notably, the performance of CCM has been found from three previous case studies to be potentially more suitable for longitudinal multi-omics analysis compared to traditional methods.

**Table 2.** Comparison of CAT Bridge with other web-based multi-omics tools.

|  | CAT<br>Bridge | OmicsAnalyst | 3omics | IntLIM | CorDiffViz |
| --- | --- | --- | --- | --- | --- |
| Pre-processing | ✓ | ✓ | × | × | × |
| Visual analytics | ✓ | ✓ | ✓ | ✓ | ✓ |
| Cross-omics association | ✓ | × | ✓ | ✓ | ✓ |
| Cross-platform | ✓ | × | × | ✓ | ✓ |
| Longitudinal study | ✓ | × | × | × | × |
| Third-party database<br>connectivity | × | × | × | × | × |
| Enrichment analysis | × | × | ✓ | × | × |
| AI agent | ✓ | × | × | × | × |

Symbols indicating feature assessments: '✓' denotes presence, '×' signifies absence. OmicsAnalyst [41]; 3omics [42]; IntLIM [43]; CorDiffViz [44].

## Discussion

In recent years, there has been a surge in multi-omics research. A critical aspect often overlooked in such studies is the unique nature of the longitudinal experimental design. Longitudinal omics analysis is particularly important in research on the developmental cycle of plants and investigations related to chronic diseases and aging [45–48]. However, many studies tend to use generic methodologies for analysis [11, 12]. This may inadvertently miss key discoveries. CAT Bridge provides a platform specifically for longitudinal multi-omics analysis, by drawing insights from disciplines where time series data is more prevalent and benchmarking them with data. Through the gene-metabolite causality and correlation modeling method, combined with visualization tools and AI assistance, researchers can more quickly identify putative genes for experimental validation. We believe that discovering associated gene-metabolite pairs will have practical applications in many fields. For example, in metabolic engineering, a single metabolite is often regulated by multiple genes, by comparing the association strength, one can infer which genes may play a dominant role under certain conditions, facilitating the development of targeted metabolic engineering strategies.

Furthermore, in horticulture and breeding, identifying genes with causal relationships to key natural products, particularly in non-model organisms, can significantly aid in molecular breeding efforts, potentially leading to the development of new crop varieties with enhanced yields or desired characteristics.

In three case studies, CCM showed its superiority compared to other methods, which is probably due to its modeling capability of dynamical systems. Thus, it can better capture the complex non-linear interactions within biological systems [21]. We advocate for modeling cause-and-effect relationships in longitudinal omics analyses, instead of more widely used Pearson or Spearman Correlations. However, this doesn’t mean that CCM is always appropriate. Factors such as sampling intervals and the number of samples also need to be considered. More precise methods for estimating causality, as well as post-processing for vector represent gene-metabolite pairs are both required to explore and validate by using more data. We tested the performance of CAT Bridge on the gene-to-metabolite and metabolite-to-gene tasks in this research, since CAT Bridge is derived by integrating algorithms originating from different fields and its theoretical basis suggests it has generalizability, we believe that it can be used for investigating associations between other different molecular levels (such as gene to protein). Nonetheless, further validating its performance across these different interactions is necessary for the conclusion due to the variations in complexity and regulatory timing at different molecular levels. Additionally, LLM-based AI agents have shown a wide range of applications in various fields, but the hallucination of knowledge deficiency remains an issue that can mislead users [49, 50]. Although we have enhanced the credibility of the AI agent response via appropriate prompt and low temperature [50–52], this cannot be regarded as a substitute for professional knowledge and experimental verification, but merely serves as a starting point for verification, the definitive conclusions still require manual assessment by researchers.

Aside from computational methods, the reliability of analytical results is also influenced by experimental design and data acquisition methods. Increasing the number of sampling time points and setting a reasonable interval between them can enhance the credibility of the results.

On the data acquisition front, it is recommended to annotate the transcriptome with an updated, high-quality reference genome, and employ advanced metabolomics techniques such as chemical isotope labeling liquid (CIL) LC-MS [53] to ensure a high coverage and more accurate relative quantification of metabolome.

Future work will involve determining response times between molecules at different levels, visualizing these lagged non-linear relationships, and constructing molecular networks based on temporal causality and metabolic pathways.

## Availability of source code and requirements

Project name: CAT Bridge

Project home page: https://catbridge.work [54]

GitHub page: https://github.com/Bowen999/CAT-Bridge

Operating system(s): Platform independent

Programming language: Python, R, HTM, CSS, JavaScript

License: CC0 1.0 Public Domain Dedication

Biotools: cat_bridge

RRID: SCR_025410

## Additional Files

**Supplementary Text 1.** Extended experimental procedure

**Supplementary Table S1.** Heuristic ranking results of Case Study 1

**Supplementary Table S2** Heuristic ranking results of Case Study 3

**Supplementary Fig. S1.** PCA plot from the test results of Case Study 1

(A) PCA plot of transcriptomics. (B) PCA plot of Metabolomics. (C) PCA plot of integrated multi-omics

**Supplementary Fig. S2.** Heatmap from the test results of Case Study 1

(A) Heatmap of transcriptomics. (B) Heatmap of Metabolomics.

**Supplementary Fig. S3.** Volcano plot (peak vs decline points) from the test result of Case Study 1

**Supplementary Fig. S4.** VIP plot of gene from the test result of Case Study 1

## Author Contribution

B Yang: Conceptualization, Software, Methodology, Visualization, Investigation, Writing— original draft. T Meng: Software. X Wang: Visualization. J Li: Investigation. S Zhao: Writing—review & editing. Y Wang: Methodology, Writing—review & editing. S Yi: Investigation. Y Zhou: Software. Y Zhang: Software. L Li: Supervision. L Guo: Conceptualization, Writing—review & editing, Supervision, Funding acquisition. All authors have read and agreed to the published version of the manuscript.

## Acknowledgement

We would like to thank the Bioinformatics Platform at Peking University Institute of Advanced Agricultural Sciences for providing the high-performance computing resources.

## Data availability

The CAT Bridge web server [54] is open and free for all users and there is no login requirement. The source code used for CAT Bridge is available on FigShare [55]. An archival copy of the code is available via Software Heritage [56].

The sequencing data for case study 1 is available in the Small Read Archive (SRA) [57] under the BioProject accession code PRJNA1030882, and the metabolome data for this case study has been deposited at the Metabolomics Workbench [58] with the study ID ST003172. The sequencing data of case study 2 was obtained from Gene Expression Omnibus (GEO) [59], with the accession number GSE85358. The sequencing data of case study 3 was obtained from GEO with the accession code GSE131992, and metabolomics data was sourced from the MetaboLights database [60] with the accession code MTBLS1044.

## References

1. Wörheide MA, Krumsiek J, Kastenmüller G and Arnold M. Multi-omics integration in biomedical research - A metabolomics-centric review. Anal Chim Acta. 2021;1141:144–62. doi:10.1016/j.aca.2020.10.038.

2. Hasin Y, Seldin M and Lusis A. Multi-omics approaches to disease. Genome Biology. 2017;18 1:83. doi:10.1186/s13059-017-1215-1.

3. Subramanian I, Verma S, Kumar S, Jere A and Anamika K. Multi-omics Data Integration, Interpretation, and Its Application. Bioinform Biol Insights. 2020;14:1177932219899051. doi:10.1177/1177932219899051.

4. Eicher T, Kinnebrew G, Patt A, Spencer K, Ying K, Ma Q, et al. Metabolomics and Multi-Omics Integration: A Survey of Computational Methods and Resources. Metabolites. 2020;10 5:202.

5. Zhou G, Ewald J and Xia J. OmicsAnalyst: a comprehensive web-based platform for visual analytics of multi-omics data. Nucleic Acids Research. 2021;49 W1:W476–W82. doi:10.1093/nar/gkab394.

6. Krassowski M, Das V, Sahu SK and Misra BB. State of the Field in Multi-Omics Research: From Computational Needs to Data Mining and Sharing. Front Genet. 2020;11:610798. doi:10.3389/fgene.2020.610798.

7. Athieniti E and Spyrou GM. A guide to multi-omics data collection and integration for translational medicine. Computational and Structural Biotechnology Journal. 2023;21:134–49. 10.1016/j.csbj.2022.11.050.

8. Albaradei S, Thafar M, Alsaedi A, Van Neste C, Gojobori T, Essack M, et al. Machine learning and deep learning methods that use omics data for metastasis prediction. Comput Struct Biotechnol J. 2021;19:5008–18. doi:10.1016/j.csbj.2021.09.001.

9. Cavill R, Jennen D, Kleinjans J and Briedé JJ. Transcriptomic and metabolomic data integration. Briefings in Bioinformatics. 2015;17 5:891-901. doi:10.1093/bib/bbv090.

10. Chong J and Xia J. Computational Approaches for Integrative Analysis of the Metabolome and Microbiome. Metabolites. 2017;7 4:62.

11. Li Y, Chen Y, Zhou L, You S, Deng H, Chen Y, et al. MicroTom metabolic network: rewiring tomato metabolic regulatory network throughout the growth cycle. Molecular plant. 2020;13 8:1203–18.

12. Yang C, Shen S, Zhou S, Li Y, Mao Y, Zhou J, et al. Rice metabolic regulatory network spanning the entire life cycle. Molecular Plant. 2022;15 2:258–75.

13. Singh KS, van der Hooft JJJ, van Wees SCM and Medema MH. Integrative omics approaches for biosynthetic pathway discovery in plants. Natural Product Reports. 2022;39 9:1876–96. doi:10.1039/D2NP00032F.

14. Ye H, Deyle ER, Gilarranz LJ and Sugihara G. Distinguishing time-delayed causal interactions using convergent cross mapping. Scientific Reports. 2015;5 1:14750. doi:10.1038/srep14750.

15. Yuan AE and Shou W. Data-driven causal analysis of observational biological time series. Elife. 2022;11 doi:10.7554/eLife.72518.

16. Sattar N and Preiss D. Reverse Causality in Cardiovascular Epidemiological Research. Circulation. 2017;135 24:2369–72. doi:doi:10.1161/CIRCULATIONAHA.117.028307.

17. Rockwood AL, Crockett DK, Oliphant JR and Elenitoba-Johnson KS. Sequence alignment by cross-correlation. J Biomol Tech. 2005;16 4:453–8.

18. Skutkova H, Vitek M, Babula P, Kizek R and Provaznik I. Classification of genomic signals using dynamic time warping. BMC Bioinformatics. 2013;14 10:S1. doi:10.1186/1471-2105-14-S10-S1.

19. Seoane JA, Campbell C, Day IN, Casas JP and Gaunt TR. Canonical correlation analysis for gene-based pleiotropy discovery. PLoS Comput Biol. 2014;10 10:e1003876. doi:10.1371/journal.pcbi.1003876.

20. Jiang MZ, Aguet F, Ardlie K, Chen J, Cornell E, Cruz D, et al. Canonical correlation analysis for multi-omics: Application to cross-cohort analysis. PLoS Genet. 2023;19 5:e1010517. doi:10.1371/journal.pgen.1010517.

21. Yuan AE and Shou W. Data-driven causal analysis of observational biological time series. eLife. 2022;11:e72518. doi:10.7554/eLife.72518.

22. Heerah S, Molinari R, Guerrier S and Marshall-Colon A. Granger-causal testing for irregularly sampled time series with application to nitrogen signalling in Arabidopsis. Bioinformatics. 2021;37 16:2450–60. doi:10.1093/bioinformatics/btab126.

23. Stokes PA and Purdon PL. A study of problems encountered in Granger causality analysis from a neuroscience perspective. Proc Natl Acad Sci U S A. 2017;114 34:E7063–e72. doi:10.1073/pnas.1704663114.

24. Arora S, Pattwell SS, Holland EC and Bolouri H. Variability in estimated gene expression among commonly used RNA-seq pipelines. Scientific Reports. 2020;10 1:2734. doi:10.1038/s41598-020-59516-z.

25. Love MI, Huber W and Anders S. Moderated estimation of fold change and dispersion for RNA-seq data with DESeq2. Genome Biology. 2014;15 12:550. doi:10.1186/s13059-014-0550-8.

26. Ye J, Coulouris G, Zaretskaya I, Cutcutache I, Rozen S and Madden TL. Primer-BLAST: a tool to design target-specific primers for polymerase chain reaction. BMC Bioinformatics. 2012;13:134. doi:10.1186/1471-2105-13-134.

27. Cantalapiedra CP, Hernández-Plaza A, Letunic I, Bork P and Huerta-Cepas J. eggNOG-mapper v2: Functional Annotation, Orthology Assignments, and Domain Prediction at the Metagenomic Scale. Molecular Biology and Evolution. 2021;38 12:5825–9. doi:10.1093/molbev/msab293.

28. Chen S, Zhou Y, Chen Y and Gu J. fastp: an ultra-fast all-in-one FASTQ preprocessor. Bioinformatics. 2018;34 17:i884–i90. doi:10.1093/bioinformatics/bty560.

29. Kim S, Park J, Yeom SI, Kim YM, Seo E, Kim KT, et al. New reference genome sequences of hot pepper reveal the massive evolution of plant disease-resistance genes by retroduplication. Genome Biol. 2017;18 1:210. doi:10.1186/s13059-017-1341-9.

30. Dobin A, Davis CA, Schlesinger F, Drenkow J, Zaleski C, Jha S, et al. STAR: ultrafast universal RNA-seq aligner. Bioinformatics. 2013;29 1:15–21. doi:10.1093/bioinformatics/bts635.

31. Pertea M, Pertea GM, Antonescu CM, Chang T-C, Mendell JT and Salzberg SL. StringTie enables improved reconstruction of a transcriptome from RNA-seq reads. Nature Biotechnology. 2015;33 3:290–5. doi:10.1038/nbt.3122.

32. Kuehne A, Hildebrand J, Soehle J, Wenck H, Terstegen L, Gallinat S, et al. An integrative metabolomics and transcriptomics study to identify metabolic alterations in aged skin of humans in vivo. BMC Genomics. 2017;18 1:169. doi:10.1186/s12864-017-3547-3.

33. Lempp M, Farke N, Kuntz M, Freibert SA, Lill R and Link H. Systematic identification of metabolites controlling gene expression in E. coli. Nature Communications. 2019;10 1:4463. doi:10.1038/s41467-019-12474-1.

34. Fattori V, Hohmann MS, Rossaneis AC, Pinho-Ribeiro FA and Verri WA. Capsaicin: Current Understanding of Its Mechanisms and Therapy of Pain and Other Pre-Clinical and Clinical Uses. Molecules. 2016;21 7 doi:10.3390/molecules21070844.

35. Kim S, Park M, Yeom S-I, Kim Y-M, Lee JM, Lee H-A, et al. Genome sequence of the hot pepper provides insights into the evolution of pungency in Capsicum species. Nature Genetics. 2014;46 3:270–8. doi:10.1038/ng.2877.

36. Chiang C-J, Ho Y-J, Hu M-C and Chao Y-P. Rewiring of glycerol metabolism in Escherichia coli for effective production of recombinant proteins. Biotechnology for Biofuels. 2020;13 1:205. doi:10.1186/s13068-020-01848-z.

37. Kuo TC, Tian TF and Tseng YJ. 3Omics: a web-based systems biology tool for analysis, integration and visualization of human transcriptomic, proteomic and metabolomic data. BMC Syst Biol. 2013;7:64. doi:10.1186/1752-0509-7-64.

38. Siddiqui JK, Baskin E, Liu M, Cantemir-Stone CZ, Zhang B, Bonneville R, et al. IntLIM: integration using linear models of metabolomics and gene expression data. BMC Bioinformatics. 2018;19 1:81. doi:10.1186/s12859-018-2085-6.

39. Eicher T, Spencer KD, Siddiqui JK, Machiraju R and Mathé EA. IntLIM 2.0: identifying multi-omic relationships dependent on discrete or continuous phenotypic measurements. Bioinformatics Advances. 2023;3 1 doi:10.1093/bioadv/vbad009.

40. Yu S, Drton M, Promislow DEL and Shojaie A. CorDiffViz: an R package for visualizing multi-omics differential correlation networks. BMC Bioinformatics. 2021;22 1:486. doi:10.1186/s12859-021-04383-2.

41. OmicsAnalyst. https://www.omicsanalyst.ca. Accessed 19 May 2024.

42. 3omics. https://3omics.cmdm.tw. Accessed 19 May 2024.

43. IntLIM. https://intlim.ncats.io. Accessed 19 May 2024.

44. CorDiffViz. https://diffcornet.github.io/CorDiffViz/demo.html. Accessed 19 May 2024.

45. Kudryashova KS, Burka K, Kulaga AY, Vorobyeva NS and Kennedy BK. Aging Biomarkers: From Functional Tests to Multi-Omics Approaches. Proteomics. 2020;20 5–6:e1900408. doi:10.1002/pmic.201900408.

46. Cellerino A and Ori A. What have we learned on aging from omics studies? Seminars in Cell & Developmental Biology. 2017;70:177–89. 10.1016/j.semcdb.2017.06.012.

47. Allegri M, Gregori MD, Minella CE, Klersy C, Wang W, Sim M, et al. ‘Omics’ biomarkers associated with chronic low back pain: protocol of a retrospective longitudinal study. BMJ Open. 2016;6 10:e012070. doi:10.1136/bmjopen-2016-012070.

48. Mars RAT, Yang Y, Ward T, Houtti M, Priya S, Lekatz HR, et al. Longitudinal Multi-omics Reveals Subset-Specific Mechanisms Underlying Irritable Bowel Syndrome. Cell. 2020;182 6:1460–73.e17. 10.1016/j.cell.2020.08.007.

49. Mittelstadt B, Wachter S and Russell C. To protect science, we must use LLMs as zero-shot translators. Nature Human Behaviour. 2023;7 11:1830–2. doi:10.1038/s41562-023-01744-0.

50. Rosoł M, Gąsior JS, Łaba J, Korzeniewski K and Młyńczak M. Evaluation of the performance of GPT-3.5 and GPT-4 on the Polish Medical Final Examination. Scientific Reports. 2023;13 1:20512. doi:10.1038/s41598-023-46995-z.

51. Antaki F, Milad D, Chia MA, Giguère C-É, Touma S, El-Khoury J, et al. Capabilities of GPT-4 in ophthalmology: an analysis of model entropy and progress towards human-level medical question answering. British Journal of Ophthalmology. 2023:bjo-2023-324438. doi:10.1136/bjo-2023-324438.

52. Miotto M, Rossberg N and Kleinberg B. Who is GPT-3? An exploration of personality, values and demographics. arXiv preprint arXiv:220914338. 2022.

53. Zhao S, Li H, Han W, Chan W and Li L. Metabolomic Coverage of Chemical-Group-Submetabolome Analysis: Group Classification and Four-Channel Chemical Isotope Labeling LC-MS. Anal Chem. 2019;91 18:12108–15. doi:10.1021/acs.analchem.9b03431.

54. CAT Bridge (Compounds And Transcripts Bridge). https://catbridge.work. Accessed 19 Mar 2024.

55. Yang, Bowen. CAT Bridge.zip. figshare. Software. 2024. 10.6084/m9.figshare25044854.v3.

56. Yang, Bowen. CAT Bridge. Software Heritage archive. 2024. https://archive.softwareheritage.org/swh:1:rev:69cf3a9cc195870d8e012a3b32e2c60a9e1a89e6;origin=https://github.com/Bowen999/CAT-Bridge;visit=swh:1:snp:176c4df277b17d55d315228e446158b5b3a0674b.

57. Sequence Read Archive. http://www.ncbi.nlm.nih.gov/sra. Accessed 19 May 2024.

58. Metabolomics Workbench. https://www.metabolomicsworkbench.org. Accessed 19 May 2024.

59. Gene Expression Omnibus. https://www.ncbi.nlm.nih.gov/geo/. Accessed 19 May 2024.

60. MetaboLights. https://www.ebi.ac.uk/metabolights/. Accessed 19 May 2024.

